# Dynamics of chromatin accessibility during human first-trimester neurodevelopment

**DOI:** 10.1101/2023.08.18.553878

**Authors:** Camiel C.A. Mannens, Lijuan Hu, Peter Lönnerberg, Marijn Schipper, Caleb Reagor, Xiaofei Li, Xiaoling He, Roger A. Barker, Erik Sundström, Danielle Posthuma, Sten Linnarsson

## Abstract

The human brain is capable of highly complex functions that develops through a tightly organized cascade of patterning events, expressed transcription factors and changes in chromatin accessibility. While extensive datasets exist describing gene expression across the developing brain with single-cell resolution, similar atlases of chromatin accessibility have been primarily focused on the forebrain. Here, we focus on the chromatin landscape and paired gene expression across the developing human brain to provide a comprehensive single cell atlas during the first trimester (6 - 13 post-conceptional weeks). We identified 135 clusters across half a million nuclei and using the multiomic measurements linked candidate *cis-*regulatory elements (cCREs) to gene expression. We found an increase in the number of accessible regions driven both by age and neuronal differentiation. Using a convolutional neural network we identified putative functional TF-binding sites in enhancers characterizing neuronal subtypes and we applied this model to cCREs upstream of *ESRRB* to elucidate its activation mechanism. Finally, by linking disease-associated SNPs to cCREs we validated putative pathogenic mechanisms in several diseases and identified midbrain-derived GABAergic neurons as being the most vulnerable to major depressive disorder related mutations. Together, our findings provide a higher degree of detail to some key gene regulatory mechanisms underlying the emergence of cell types during the first trimester. We anticipate this resource to be a valuable reference for future studies related to human neurodevelopment, such as identifying cell type specific enhancers that can be used for highly specific targeting in *in vitro* models.

## Introduction

Through a tightly organized cascade of patterning, specification and differentiation events, the human brain develops into a highly complex system capable of unique cognitive abilities beyond those of other mammals. The human brain consists of more than a thousand distinct types of neurons, glia, and non-neural cell types^1^. Single-cell RNA-sequencing has enabled parallel profiling of cell types and states, revealing both regional differences and subtle variation between closely related cell types ^2–4^. Profiling the developing human brain has revealed differentiation trajectories leading to diverse neuronal and non-neuronal cell types^5^. During development, the functional architecture of the genome is constantly in flux, with changes in the expression, binding and regulation of transcription factors (TF) driving cell fate decisions. The activities of regulatory elements in development are often both cell type-specific and brief. This dynamism complicates the interpretation of Genome Wide Association Studies (GWAS) of complex neurodevelopmental disorders, because identified loci — which predominantly fall in the non-coding DNA — are equally context specific^6,7^. Previous work has mapped the regulatory landscapes of the developing human brain in organoids and induced pluripotent stem cell-derived model systems^8^, the second trimester developing cortex^9^, and in whole embryos^10^. Here we focus on the chromatin landscape across the whole developing human brain during the first trimester, a pivotal time when the brain is patterned and many neural cell types acquire their core transcriptional identities.

### Chromatin accessibility in the first trimester

We measured chromatin accessibility in the developing human brain from 6 to 13 post-conception weeks (p.c.w.; Fig. 1d) using the 10XGenomics single-cell assay of transposase-accessible chromatin (scATAC-seq^11^; 18 specimens), a combined scATAC and scRNA-seq assay (Multiome; three specimens) or both (five specimens). Each specimen was dissected into the major antero-posterior segments (Fig. 1a-c; telencephalon, diencephalon, mesencephalon, metencephalon and cerebellum). We collected relatively more nuclei from the brain stem region, which is highly complex but has been comparatively less studied than the forebrain^9,10^. After removing low-quality nuclei (Methods), we collected chromatin profiles from a total of 526,094 nuclei and 76 unique biological samples (116 including technical replicates; Extended fig. 1g). 166,785 of these nuclei included gene expression profiles from Multiome sequencing.

**Figure 1:**
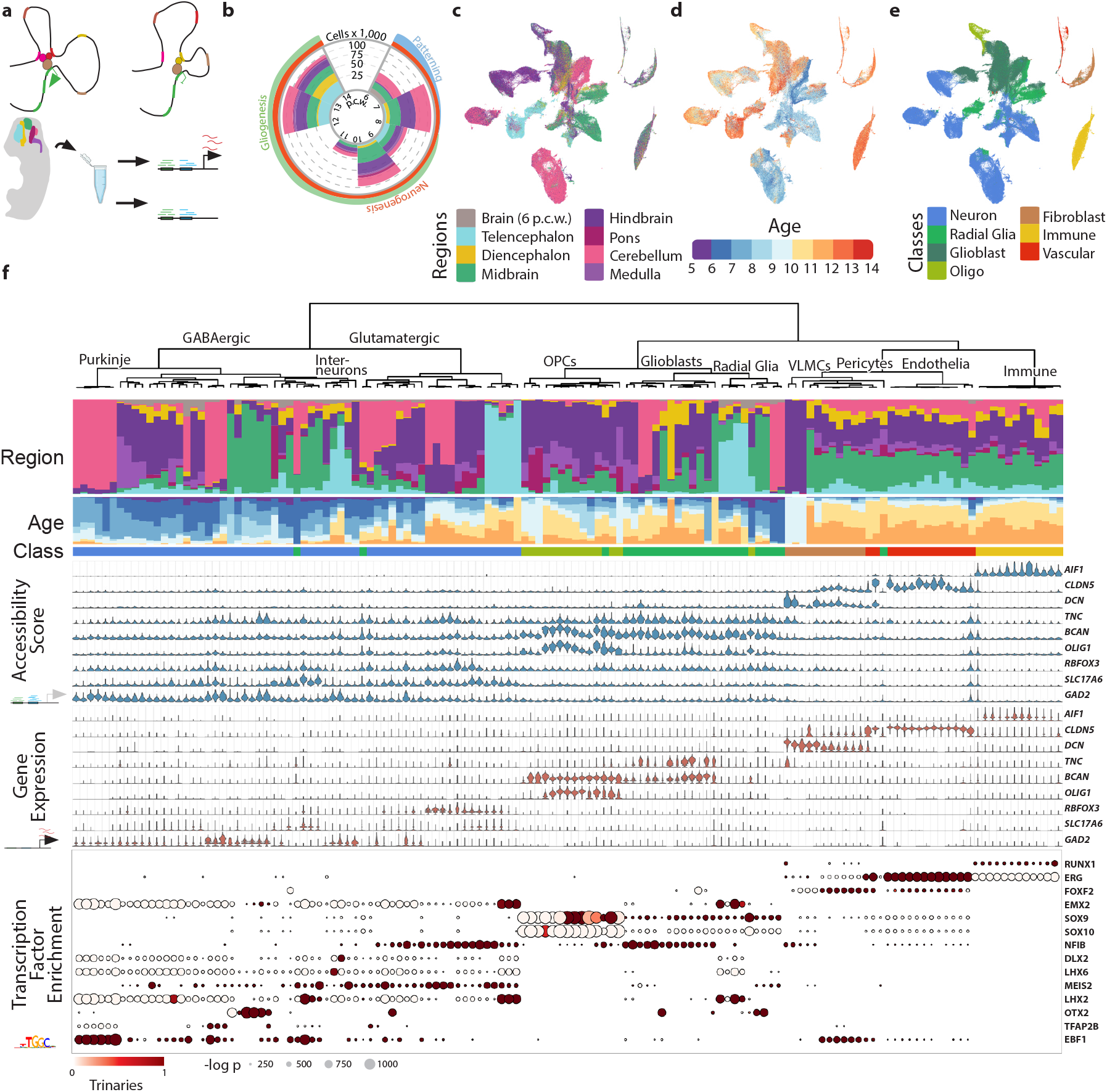
Atlas overview. A) Overview of experimental design. Dissected samples were processed using scATAC-seq and single-cell Multiome sequencing to infer gene-regulatory relationships throughout early development across the human brain. B) Collected cell counts per post-conceptual week (p.c.w.) and region. C) tSNE plot of regional identities. D) Distribution of developmental ages across the tSNE-embedding. E) tSNE plot of cell classes. F) From top to bottom. Per cluster regional distribution of cell types, (legend in C). Distribution of ages (legend in D). Assigned cell class (legend in E). Aggregated gene activity by cluster based on region co-accessibility. Gene expression of the same genes. Transcription factor motif enrichment in the top 2,000 most enriched accessible regions per cluster. Dot size represents motif enrichment, while color indicates corresponding expression of the transcription factor (trinarization score; a probabilistic score of whether a gene is expressed).

To identify the feature set of accessible regions, we applied stratified peak calling on a rough clustering of the data using 20 kb genomic bins as temporary features. This was followed by a more robust clustering based on the accessible regions using Latent Semantic Indexing and Louvain clustering. Batch correction was performed using Harmony^12^. A split-and-pool approach was then used to subcluster each cell class (Radial Glia/Glioblast, OPC, Neuron, Fibroblast, Vascular and Immune), resulting in 135 clusters (Fig. 1e-f).

About 36% of accessible regions were intergenic, while 64% were located in gene bodies or promoter regions with the majority being intronic (51%; Extended fig. 4a). When taking only the distance to the transcription start site (TSS) into account, 87% of accessible regions were marked as distal (>2 kb from the TSS) and 19,494 accessible regions (4.7%) overlapped with known TSS sites (Extended fig. 4b). Additionally, 18% of elements overlapped with a transposable element (Extended fig. 4i), compared to 22% in similar data from the adult human brain^13^.

Access to gene expression data also allowed for better identification of candidate *cis-* regulatory elements (cCREs) using a modified version of Cicero^14^. By leveraging co-occurrence of chromatin accessibility and gene expression, 106,991 predicted enhancer-gene interactions were identified for 16,267 genes and 59,069 accessible regions (henceforth, cCREs).

The dataset primarily consisted of nuclei from the neural lineage, showing strong regional identities, while non-neural clusters including endothelial cells, fibroblasts and microglia showed limited spatial identities (Extended fig. 3a). Additionally, the radial glia to glioblast ratio differed markedly between regions (Extended fig. 3), with posterior regions being enriched for glioblasts while the more anterior regions showed primarily radial glia (Extended fig. 3c). This difference in abundance is most likely the consequence of a later transition from radial glia to glioblasts^5^ in the anterior brain.

The Multiome data allowed us to impute gene expression across the dataset, providing a direct comparison between gene expression, gene accessibility and the enrichment of transcription factor binding motifs. We combined conventional motif discovery with gene expression for each cell type to limit identified transcription factor motifs to those coinciding with transcription factor expression, discarding unexpressed redundant motifs. The identified motifs included early neuronal (*EBF1*) and pan-glial markers (*SOX9*) as well as TFs with strong lineage-specific expression (e.g. *LHX6* and *DLX2* in MGE- and LGE/CGE-derived interneurons, *EMX2* in telencephalic glutamatergic neurons, *OTX2* in Midbrain GABAergic neurons; additional TFs in Fig. 1f).

We observed a significant 10% increase in the number of accessible regions along the neuronal differentiation trajectory, but no significant increase in the glioblast lineage (p < 0.05; Extended fig. 3g). This is in line with a shift towards heterochromatin that has been observed in oligodendrocytes where large numbers of neuronal and later OPC genes are silenced during differentiation^15,16^. Interestingly, the number of accessible regions also increased by age across all classes except radial glia (p < 0.001; coefficient 3,206; SE 634; t = 5.06; 6 DF; linear regression), with the newly acquired accessible regions being strongly enriched for NFI-binding sites (Fig. 2a-b). Similarly only within the radial glia and glioblast classes did we find that NFI-binding sites were most associated with more mature cell neighborhoods (Milopy^17^; Extended fig. 3d-f).

**Figure 2:**
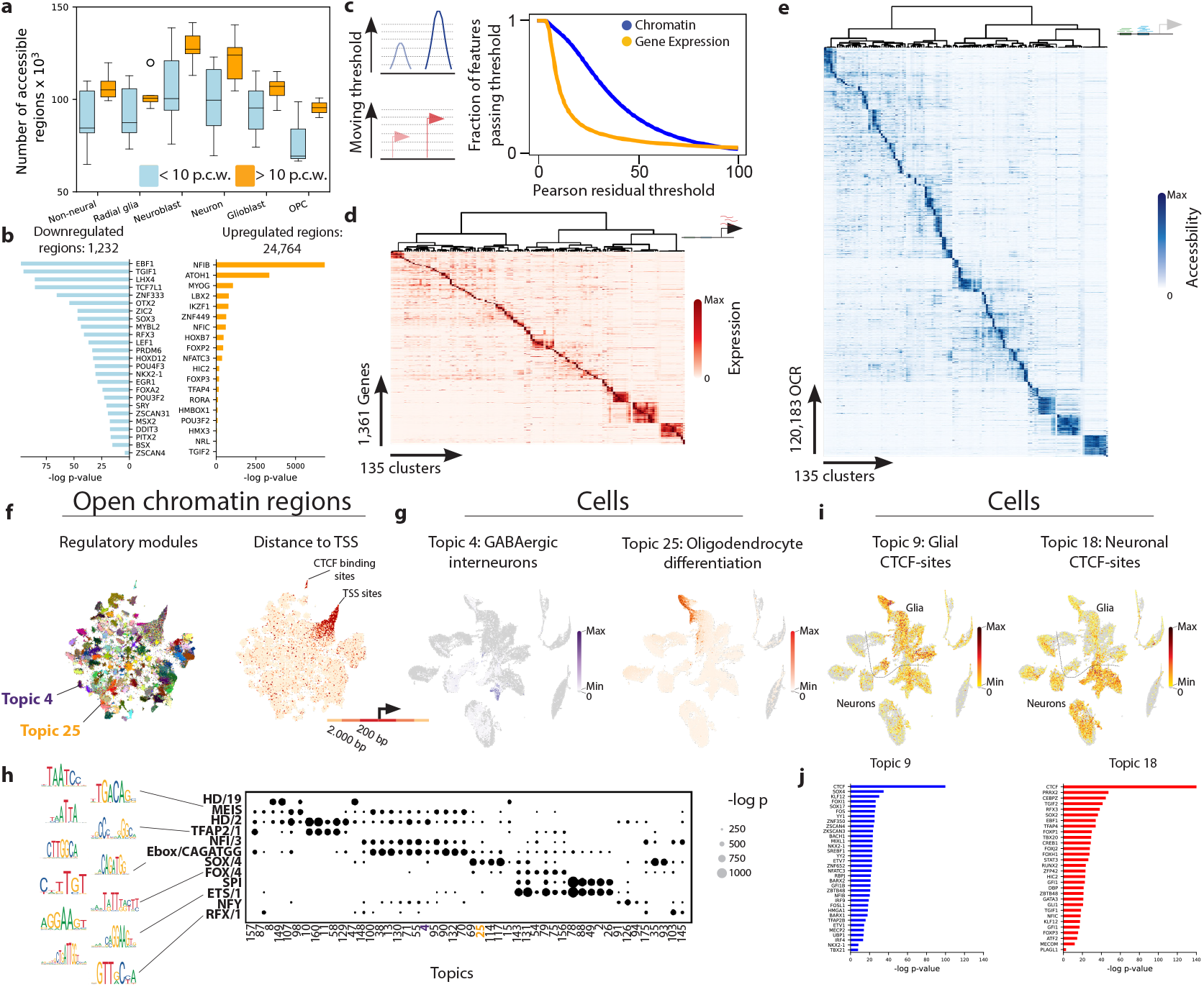
Functional Annotation of Open Chromatin Regions. A) Number of accessible regions by cell class, split between samples staged before and after 10 p.c.w. B) Transcription factor motifs in regions that are differentially accessible early or late in the dataset (p < 0.05; Benjamini Hochberg-corrected one-sided Fisher exact test). C) Enrichment comparison between the gene expression and chromatin accessibility components of the dataset. A moving threshold was used to identify the fraction of features enriched in at least one cluster at different levels of stringency. D) Selection of marker gene expression. E) Accessible marker regions, limited to top 2,000 per cluster. F) tSNE plot where individual dots represent accessible regions and are colored based on highest scoring region-topic. A strong enrichment TSS and promoter regions in the top right of the second plot represent constitutively active elements. G) tSNE plot of nuclei showing enrichment of two Gene Ontology signatures that were enriched in topic 4 and 25 (shown in F), identified using GREAT. H) Arche-motif enrichment for a subset of topics, where dot size represents enrichment. The arche-motifs from top to bottom contain the following transcription factors (non-exhaustive): HD/19: OTX1/2, MEIS: MEIS2/3, HD/2: HOXA2/LBX2, TFAP2/1: TFAP2A/B, NFI/3: NFIA/C, Ebox/CAGATGG: PTF1A/NEUROD1/2/ATOH1, SOX/4: SOX4/10, FOX/4: FOXA1/2/FOXP2, SPI: SPI1/SPIB/C, ETS/1: ELF1/3/5/GABPA, NFY: NFYA/B/C, RFX/1: RFX1/2/3/4. I) tSNE plots of topic 9 and 18. Each of these was enriched for CTCF-binding sites and plotted together on the region tSNE (also shown in F). J) Enriched transcription factor motifs in topic 9 and 18.

### Cis-Regulatory elements predict gene expression and Transcription Factor specificity

We next asked how cell type specificity compared between chromatin accessibility and gene expression. We used the variance between the cluster level Pearson residuals as a measure of specificity. For most marker genes, gene expression was more specific than the sum of linked accessible regions (Extended fig. 6a). In contrast, individual accessible marker regions were generally more cell type-specific than marker genes (Fig. 2c). As a consequence, we found 1,361 marker genes, but 120,183 marker regions (Fig. 2d-e). Thus cCREs discovered here provide a rich source of regulatory elements with precise cell type- state- and temporal resolution during brain development.

We next assessed the region specificity of accessible regions by comparing them to known functional central nervous system enhancers from the VISTA developmental enhancers database^18^. Nearly all of the VISTA enhancers overlapped with accessible regions in our data (96% overlapping feature set; 39% intergenic, 53% intronic, 4% promoter). Many VISTA enhancers are specific to either the forebrain, midbrain or hindbrain, and these showed a similar pattern of activity in the scATAC-seq dataset (Extended fig. 4c). In many cases these enhancers were only accessible in more specific cellular lineages like hindbrain glutamatergic neurons (HS161; Extended fig. 4d), immature interneurons in the ganglionic eminences (HS702) or radial glia and GABAergic neurons in the midbrain (HS830).

To better understand the gene-regulatory programs underlying the dataset we identified accessible region topics using pycisTopic^19^, which uses a Latent Dirichlet Allocation model (LDA) to identify groups of accessible regions that covary and are likely to represent biological programs. Each cluster was downsampled to 1,000 nuclei and we fitted a model with 175 topics based on the point where the log-likelihood estimation and topic coherence scores reached saturation. A t-SNE embedding of the accessible regions based on the topic scores showed distinct clusters linked to individual topics (Fig. 2f), representing distinct regulatory programmes. Interestingly, in contrast to distal elements, most TSS regions were not strongly linked to individual topics and clustered together on the embedding, indicating that they were less variable and represent constitutively open promoters. A subset of promoter proximal regions clustered separately, and represented two topics of pan-neuronal and glial CTCF-binding sites (Fig. 2i-j).

We used the Genomic Regions Enrichment for Annotation Tool (GREAT) to link topics to known biological processes via the biological annotation of nearby genes (Extended data 3). For example, Topic 4 and 25 were enriched for genes relevant to GABAergic interneuron identity and oligodendrocyte differentiation respectively. When scoring the associated signatures (accessible regions in the topic related to the pathway) clear enrichments in the immature interneuron and oligodendrocyte precursor populations could be identified (Fig. 2g), respectively. Since individual topics reflected region accessibility only and not gene expression, we identified enriched TF motifs for each topic and reduced them to a set of archemotifs^20^. This prevented the prioritization of false positive motifs based on the similarity of the binding motif within TF families. Indeed, Topic 4 was enriched for the MEIS (i.a. *MEIS2*), HD/2 (i.a. *DLX2/5*), Ebox/CAGATGG (i.a. *NEUROD1*) and NFI (i.a. *NFIA/B/X*) archemotifs, while Topic 25 was primarily enriched for the SOX/4 (i.a. *SOX10*) archemotif (Fig. 2h).

### Enhancer logic in neuronal specification

While topic modeling can be a useful tool to understand the activity of accessible regions, it does not offer any explanations as to the underlying logic that drives activity of regions between cellular lineages. To better understand the syntax of regulatory elements that differentiate neuronal lineages we trained a convolutional neural network (CNN) to predict cell-type identity based on sequence composition^21,22^. We focused on five large, well-sampled clades: GABAergic neurons from the midbrain, glutamatergic neurons from the hindbrain and telencephalon, and granule and purkinje neurons from the cerebellum. The model consisted of four convolutional layers followed by two dense layers and was able to predict the correct class with an average ROC AUC score (Receiver Operating Characteristic Area Under Curve) of 0.92 across the classes (Fig. 3a-b). We determined the contribution of each nucleotide in the target sequences towards the prediction using DeepExplainer^23^, and identified short motifs with high predictive power (seqlets) using TF-MoDisCo^24^. In this way, we discovered on average 6 seqlets accessible region, and 84% of the selected regions were associated with at least one seqlet (online data). We identified *MEIS1* and *ATOH1* as key regulators in both hindbrain glutamatergic neurons and cerebellar granule neurons (Fig. 3c) even though *ATOH1* expression could only be detected in a subset of the nuclei in those populations (Fig. 3d). Telencephalic glutamatergic neurons on the other hand were quite distinct from their posterior counterparts and were characterized by *LHX2* and *BHLHE22* motifs. For the GABAergic neurons the *GATA2* motif was only observed in midbrain neurons, while the *OTX2* motif could also be seen in the Purkinje neurons where the gene is not expressed. This most likely represents *DMBX1*, a TF from the same family that is expressed in Purkinje progenitor cells both in this dataset and a previously published human neurodevelopment scRNA-seq dataset^5^ (Extended fig. 7a). Both populations contained the motif for *TFAP2B* and *LHX1/5*, with *LHX1* only being minimally expressed in the Midbrain neurons.

**Figure 3:**
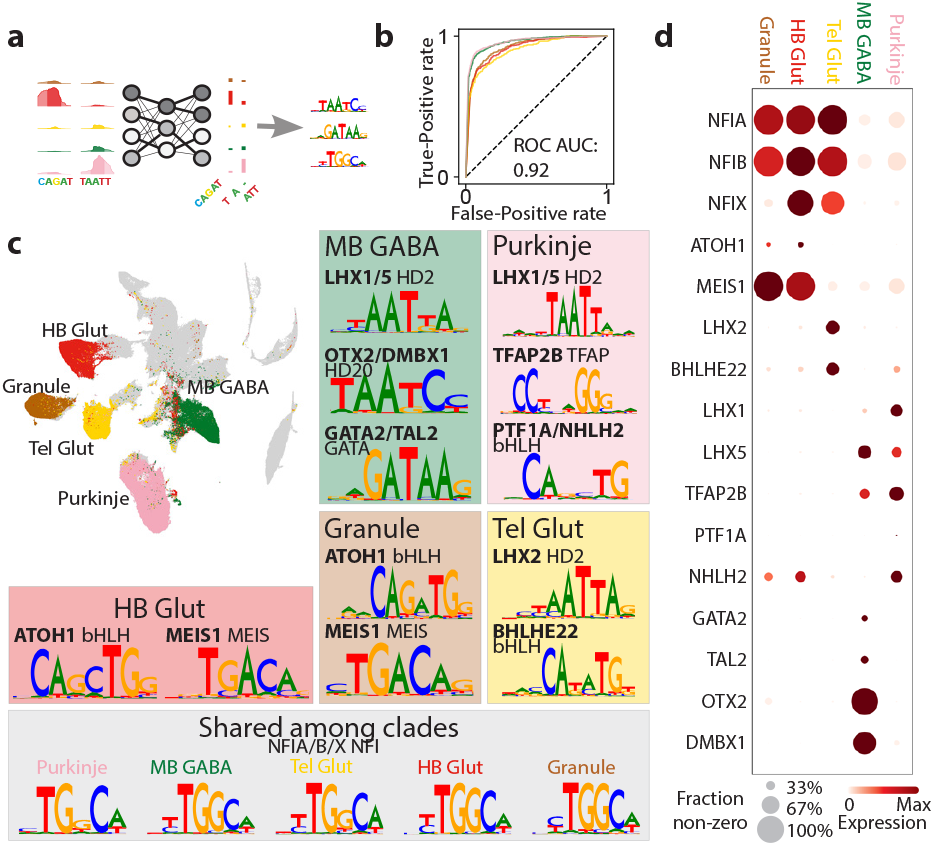
Convolutional Neural Network predicts Neuron Type from Sequences. A) Network layout. Enriched sequences for each cell type are one-hot encoded and used to train a CNN network with 4 convolutional layers and 2 dense layers. The loadings of the convolutional layers are used to calculate the contribution of each of the nucleotides in the original sequence and recurrent patterns can be clustered to find driving TF motifs. B) ROC curves for each of the classes and mean area under curve score (ROC AUC) off all classes. In a random classification model the ROC AUC is 0.5 C) t-SNE embedding showing the selected classes. For each of the classes a subset of the characteristic motifs is shown. The identity of motifs was determined by the binding motif and expression in the corresponding class. D) Dot plot representing expression of each of the transcription factors in C. The size represents the fraction of non-zero cells and the color the expression level.

### Gene regulatory dynamics in Purkinje Neuron development

The CNN did not provide a temporal order of what stages of the cell trajectory these TFs are active in. To further investigate the relationship between TF expression and cCRE accessibility we focused on the Purkinje lineage, which was well sampled in our dataset. Purkinje neurons are born in the ventricular zone of the hindbrain from *PTF1A*^+^ progenitors. From there they migrate into the developing cerebellum forming a characteristic layer of large, arborated neurons. We fitted a pseudotime trajectory to the 71,947 nuclei of the Purkinje lineage (Fig. 4a) and identified differentially expressed genes across the lineage (Fig. 4d). We next applied DELAY^25^, a different convolutional neural network method that exploits the temporal shift between the expression of transcription factors and their targets in single-cell lineages in combination with ChIP-seq derived TF binding site information to estimate gene regulatory networks (GRNs). This revealed a network of 148 TFs co-regulating each other during Purkinje cell differentiation (Fig. 4b; Extended data 4).

**Figure 4:**
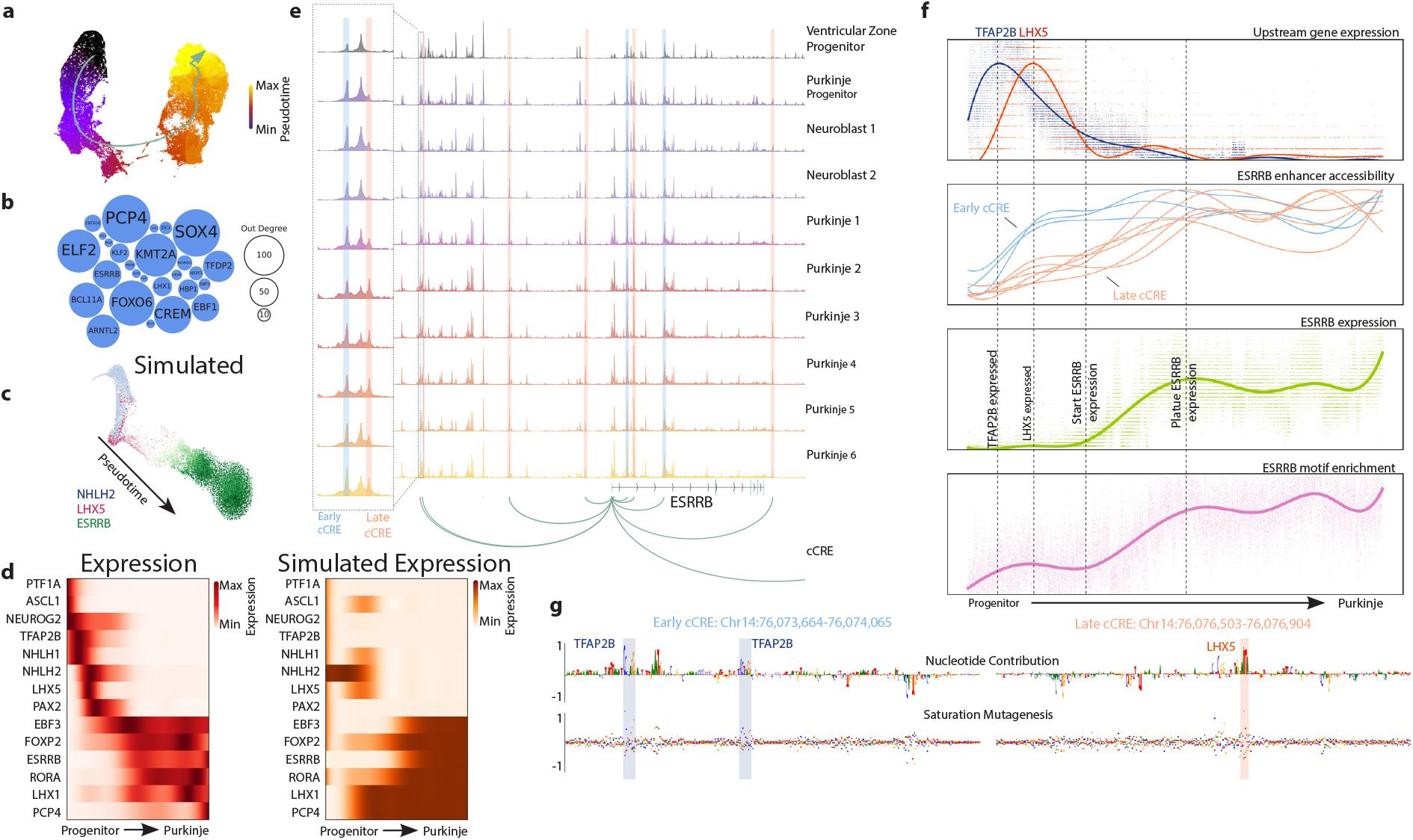
Gene Regulatory Dynamics in Purkinje Neurons. A) t-SNE embedding of Purkinje Neurons showing pseudotime and the differentiation trajectory. B) Transcription factors involved in the Gene Regulatory Network as identified using DELAY. The nodes are sized by centrality. C) t-SNE of nuclei simulated from the DELAY network using boolODE. Color represents the expression of *NHLH2*, *LHX5* and *ESRRB.* D) Heatmaps of Purkinje marker genes expressed along the pseudotime trajectory and the corresponding prediction based on the DELAY network. E) Chromatin Accessibility landscape around the *ESRRB* gene. 9 cCRE were identified to regulate *ESSRB* expression. Two are highlighted that occur close to each other but open up at different stages in differentiation. F) Trend lines of important factors in *ESSRB* gene progression. The vertical lines mark important events. From top to bottom: Expression of *LHX5* and *TFAP2B*, Accessibility of cCREs regulating *ESRRB*, Expression of the *ESRRB* gene and enrichment of the ESRRB binding-site in target peaks. G) The contribution scores of two cCRE regulating *ESRRB* expression (an early and a late cCRE), title colors correspond to the marked peaks in E. The early example contains two binding sites for TFAP2B, while the late cCRE contains a LHX5-binding site.

We used the inferred GRN to computationally model single cells, recapitulating *in silico* the expression dynamics of transcription factors along the trajectory (Fig. 4c-d). One of the central transcription factors in the network dynamics was *ESRRB*, an estrogen-related nuclear receptor TF that in the cerebellum is expressed uniquely in Purkinje neurons. Expression of *ESRRB* was preceded by a series of other transcription factors (*PTF1A*, *ASCL1* and *NEUROG2* in the progenitor phase; *NHLH1/2*, *TFAP2B*, *LHX5* and *PAX2* in the neuroblast phase) and itself preceded the expression of later Purkinje markers like *PCP4*. We identified nine cCREs linked to *ESRRB*, which showed two distinct activation patterns, early and late (Fig. 4e-f). Using the CNN we had previously trained to distinguish neuronal cell types we then identified the nucleotides driving the Purkinje lineage identity in these two groups of cCREs. We found multiple *TFAP2B* binding motifs in the early cCREs, and an increase of *LHX5* binding motifs in the late cCREs (Fig. 4g). Finally, once ESRRB was expressed, we observed increased accessibility at its downstream binding sites elsewhere in the genome (Fig. 4f). The activation of *ESRRB* can thus be seen as a two step process where the gene is first poised for expression by *TFAP2B*, after which *LHX5* binds the late cCREs and *ESRRB* expression is induced, leading eventually to the activation of *ESRRB* target genes. Our dataset provides rich resources, RNA expression for every TF (online data), predicted cCREs and their activities (online data), and predicted seqlets for every accessible region included in training the CNN (online data) to explore similar regulatory processes for many other genes and lineages.

### Chromatin accessibility and GWAS polymorphisms predict cellular targets in neuropsychiatric disorders

Mutations in non-coding gene-regulatory regions have been implicated in numerous psychiatric disorders ^26,27^. In many instances these non-coding regions are primarily active during a limited temporal window in selective cell types, which makes it difficult to identify the affected developmental processes^28^. Chromatin accessibility atlases with single-cell resolution spanning across multiple developmental time points can thus be an important tool in the identification of cell type-specific vulnerabilities in complex trait disorders by providing increased selectivity^9,13^. To identify if any of the cell types in our dataset were selectively vulnerable to mutations associated with psychiatric disorders we curated a large set of phenotypes from the UK Biobank^29^ as well as GWAS results from 11 psychiatric phenotypes^30–40^. We used stratified linkage disequilibrium-score regression to identify cell types for which the phenotype was enriched for SNPs within the corresponding cell-type specific accessible regions^41^. As all the accessible regions within our dataset are of brain tissue during early development, we wanted to ensure that cell-type enrichments for a phenotype remained significant when conditioned on other life-stages and tissues. We therefore added accessible regions identified throughout development^10^ and adulthood^42^ to the background dataset to correct for our fetal neural-focused selection of features. We found the expected associations for many of the non-neural cell types (Extended fig. 8) and a number of significant enrichments for the psychiatric phenotypes in neuronal subtypes. After correcting for multiple testing (Bonferroni or false discovery rate; Fig. 5a), no significant enrichments were found for Tourette’s syndrome, obsessive compulsive disorder, bipolar disorder, alcohol use disorder or Alzheimer’s disease, although interestingly we did see lower uncorrected P-values (Extended fig. 9; 0.003 < P < 0.01; methods) in all immune cells for Alzheimer’s disease compared to the neural cell types (all p > 0.1; methods) which agrees with previous findings linking SNPs to immune genes^43^.

**Figure 5:**
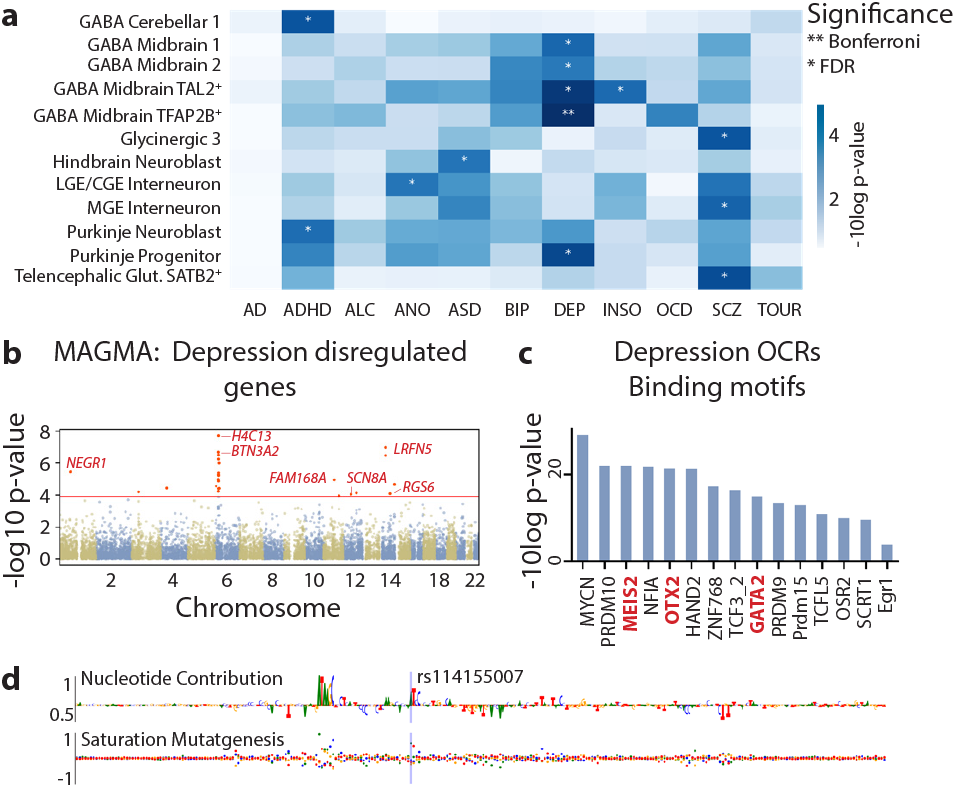
Enrichment of Psychiatric SNPs in first trimester Central Nervous System Cell Types. A) -log10 p-values for Neuropsychiatric phenotypes included in the analysis. Asterisks indicate significance level (FDR or Bonferroni). Only cell types reaching significance in one of the phenotypes are plotted. B) MAGMA -log10 p-values for gene-associated cCREs with the Benjamini-Hochberg alpha set at 1.2 x 10^4^. C) Enriched TF-binding motifs in the MDD-associated SNPs passing Bonferroni correction. MEIS2, OTX2 and GATA2 are transcription factors strongly associated with midbrain inhibitory neurons. D) The contribution scores of region chr6:28,885,244-28,885,645, which contains rs114155007, one of the SNPs associated with major depressive disorder.

Several disorders showed associations that agree with known disease biology. Schizophrenia was associated with cortical MGE-derived interneurons and *SATB2*^+^ telencephalic excitatory neurons supporting a cortical developmental origin of the disease^38^. ADHD was associated with immature GABAergic neurons and Purkinje neuroblasts in the cerebellum, which might be related to the structural abnormalities in the cerebellum often observed in ADHD patients^44^. Anorexia nervosa was associated with LGE- and CGE-derived interneurons, in agreement with known eating-disorder associated SNPs in GABAergic receptors^45^. Autism spectrum disorder (ASD) was associated with neuroblasts from the hindbrain, supporting the brainstem hypothesis of ASD^46^. For insomnia, *TAL2^+^*GABAergic neurons in the midbrain were implicated, in line with the reported role of such neurons in the reticular formation of the ventral midbrain in wakefulness^47,48^.

The strongest associations, however, were those observed between midbrain-derived GABAergic neurons (multiple groups) and major depressive disorder (MDD), which we validated in a second cohort^49^ (Extended data 7). The involvement of GABAergic neurons in MDD is well established^50^, but often attributed to cortical interneurons for which we found no significant associations. Midbrain GABAergic neurons however, are also known to be involved in the regulation of reward behavior and stress^51^, two systems known to be disturbed in MDD. Moreover, a subset of these *SOX14^+^* midbrain-derived neurons also migrate to the thalamus and pons^1^, suggesting a broader effect from these mutations. The overlap between MDD and insomnia in *TAL2^+^* midbrain GABAergic neurons is also notable as the two disorders have high comorbidity^52^.

To better understand the association between MDD and midbrain GABAergic neurons, we used cCREs to identify target genes in MDD. We pooled the set of cCREs linked to each individual gene and used MAGMA^53^ to identify genes significantly associated with MDD. This yielded 25 associated genes consisting mostly of known MDD genes like *NEGR1*, *BTN3A2*, *LRFN5* and *SCN8A*, as well as a number of histone genes located in the same locus as *BTN3A2* (Fig. 5b; *H2AC13*, *H2AC15*, *H2BC14*, H2BC15 and *H4C13*). While many of these genes were expressed in midbrain GABAergic neurons, none of them were specific to them. Conversely, regions significantly associated with MDD were enriched for the *MEIS2*, *OTX2* and *GATA2* binding motifs, indicating a midbrain GABAergic identity (Fig. 5c), with 22 of the 114 significant MDD regions also containing CNN-predicted *OTX2* binding sites and 36 containing predicted *GATA2* binding sites. We therefore examined individual accessible regions and the predicted nucleotide contributions to the midbrain GABAergic fate (DeepExplainer scores). For most MDD-associated SNPs we did not find immediately interpretable overlaps, however, rs114155007 directly overlapped with the *OTX2* binding site (Fig. 5d). These findings suggest that some broadly expressed genes associated with MDD contribute to disease only when perturbed specifically in midbrain GABergic neurons.

## Discussion

In this study we provide a high-resolution multiomic atlas of chromatin accessibility and gene expression in the first trimester human brain. We identified over a hundred thousand cell type- and region-specific developmental accessible chromatin regions, inferred cCREs, and predicted their regulatory syntax using convolutional neural network modeling. These resources enable analyses that span from developmental lineages to individual nucleotides — linking transcription factors to putative enhancers, and enhancers to their target genes — as exemplified here by our analysis of *ESRRB’*s regulation in the Purkinje neuron lineage.

Our dataset further enabled analysis of genetic association to disease. Interestingly, we found that most genes linked to MDD were not cell type specific, yet the associated accessible regions showed enriched transcription factor motifs consistent with midbrain GABAergic neurons. This suggests that dysregulation of those genes contributes to MDD only when the dysregulation affects specific midbrain cell types (but may cause other phenotypes when dysregulated in other cell types). The observation reinforces the fact that disease-associated alleles are contextual, and yield disease phenotypes mainly by their effect in specific cell types. Nonetheless, our GWAS analysis covered only a relatively early period of neurodevelopment and more complete datasets will be required to fully elucidate the genetics of complex diseases relative to brain cell types.

In conclusion, this study provides a rich resource for the study of early embryonic human neurodevelopment in the context of gene regulation and neurodevelopmental disease.

## Methods

### Data Availability

Raw sequencing data is available through from the European Genome Phenome Archive (EGAS00001007472). To facilitate ease of use of the resource the chromatin accessibility and gene expression data are browsable through the CATlas webbrowser (http://catlas.org/humanbraindev) and the convolutional neural network can be downloaded through github: https://github.com/linnarsson-lab/fetal_brain_multiomics.

### Code Availability

All code used to reproduce the figures is available through github: https://github.com/linnarsson-lab/fetal_brain_multiomics. Code to reanalyze the data is available through: https://github.com/linnarsson-lab/chromograph. The DELAY models trained on scATAC-seq data are available through https://github.com/calebclayreagor/DELAY.

### Sample collection

Human fetal samples were collected from routine termination of pregnancies at the Karolinska University Hospital, Addenbrooke’s Hospital in Cambridge and the Human Developmental Brain Resource (HDBR) following informed consent of the donors. The use of fetal samples collected from abortions was approved by the Swedish Ethical Review Authority and the National Board of Health and Welfare. In the UK, approval from National Research Ethics Service Committee East of England – Cambridge Central was obtained (Local Research Ethics Committee, 96/085). The samples were dissected by a trained embryologist into the major developmental regions (Telencephalon, Diencephalon, Mesencephalon and Metencephalon) along the anterior-posterior axis. In addition, the Cerebellum was separated from the Metencephalon and where possible the Metencephalon was divided into Medulla Oblongata and Pons. Following the dissection, the samples were transferred to ice cold Hibernate E media (ThermoFisher, A1247601) and either shipped overnight at refrigerated temperature to Sweden or processed the same day when collected at the Karolinska University Hospital. Ethical approval for the use of post-mortem human fetal tissue was provided in DNR2019-04595 and DNR2020-02074.

However, some important limitations of this study must be considered. First, as these are clinical samples the timing was variable and based on expert annotation rather than knowledge of the date of conception. In addition, due to damage incurred during collection not all regions could be collected from every sample and had to be compensated for by collecting more samples. Finally, as the samples were derived from multiple sources the time between collection and dissociation varied.

### Nuclei isolation

Tissue was gently minced using a razor blade and incubated with the Papain Dissociation System (Worthington) following the manufacturer’s recommendations (including 200 U/mL DNAse), at 37 °C for 10 minutes. Using glass pipettes, the suspension was then triturated to dissolve any remaining chunks of tissue, before being filtered through a 30 um filter (CellTrics). The cells were then washed with EBSS, concentrated (200g 5 min) and counted using a hemocytometer, after which 1×10^6^ cells were pelleted (500g 5 min) in a 2 mL LoBind Eppendorf tube and pelleted. The cell pellets were dissociated for 5 minutes on ice using 100 ul of dissociation mix (0.001% Digitonin, 0.01% Non-idet P40, 1 mM DTT, 1 U/ul RNAse inhibitor, 0.1% Tween-20, 1% BSA, 10 mM Tris-HCl, 10 mM NaCl, 3 mM MgCl_2_). When only scATAC-seq was performed no RNAse inhibitor or DTT were added to the mix. Dissociation was halted by addition of 1 ml of wash buffer, after which nuclei were pelleted again (500g 5 min) and resuspended in 1X nuclei buffer (10X Genomics) and recounted.

### Single cell sequencing

Libraries were generated using the 10X Genomics Chromium controller and Single Cell ATAC or Single Cell Multiome ATAC + Gene expression kits. Briefly, a targeted number of nuclei (5.000-10.000) was treated with a Tn5-transposase for 60 min at 37 °C to fragment the DNA and insert adapter sequences into open parts of the chromatin. The suspension was then mixed with the provided barcoding PCR mix and a gel-bead emulsion (GEM) was generated by co-encapsulating the suspension with barcoded beads in in the 10X microfluidic chip and RT-PCR was performed in a C1000 Touch Thermal Cycler (Bio-Rad) with one of two programs: 1) ATAC: 12 cycles of (5 min 72 °C, 30s 98 °C, 10s 98 °C, 30s 59 °C, 1 min 72 °C) and hold at 15 °C or 2) Multiome: 45 min 37 °C, 30 min 25 °C and hold at 4 °C. For multiome samples Quenching Agent was added to prevent the RT-PCR from continuing. Following PCR, the DNA was isolated from the droplets and cleaned up with the Cleanup mix and Silane Dynabeads. Sample indexes and P7 primers (Illumina) were ligated during library construction using the following PCR protocol: 9 or 10 cycles of (45s 98 °C, 20s 98 °C, 30s 67 °C, 20s 72 °C) and 1 min 72 °C before hold at 4 °C. SPRIselect beads were used for size selection of fragments to generate the final library. The fragment size distribution was analyzed using the Bioanalyzer high-sensitivity chip to eliminate libraries that did not show the expected nuclear banding pattern. Libraries were then sequenced using the Illumina Nova-seq instrument using the recommended setting for paired-end sequencing, with the scATAC-seq and scATAC-seq (multiome) libraries in separate flow cells as pooling of them is not recommended with a target of 100.000 read pairs per cell. The multiome scRNA-seq libraries were pooled with other 10X Genomics scRNA-seq v3.1 libraries.

### 10X data processing

All samples were demultiplexed and aligned to the human genome GRCh38.p13 gencode V35 primary sequence assembly using either Cellranger-atac 2.0.0 or Cellranger-arc 2.0.0 for scATAC-seq and single-cell multiome respectively. The RNA libraries from multiome samples were aligned as described previously^5^.

### Chromograph pipeline

Chromograph is a new analysis pipeline for scATAC-seq data based on the key architecture of Cytograph 2 ^4^, which uses loom-files as the underlying data-format and is available for use in github (https://github.com/linnarsson-lab/chromograph). The results in this paper were generated using commit #9ae1434. Briefly put, chromograph provides tools to pool and split scATAC-seq data, perform clustering, balanced peak calling based on cluster partitions, identify marker peaks and enriched transcription factor motifs and enables imputation of gene expression from limited multiome data. This dataset was analyzed by first performing a primary analysis, then manually splitting it into subsets based on marker genes. These subsets were then reanalyzed, and the results are again pooled together to generate a more fine-grained dataset than the primary analysis.

### scATAC-seq QC

TSS enrichment was calculated using pyCistopic^19^ (tss window 50bp, flanking window 1,000 bp) since we noticed discernible change in some of the samples after updating Cellranger-arc. Samples with a score below 5 were discarded. For the other samples cell-by-bin matrices were generated at both 5kb and 20kb resolution with bins that overlapped with any of the ENCODE blacklist^54^ being removed. The 5kb cell-by-bin matrix was used for doublet detection using an adapted version of DoubletFinder. Briefly put, nuclei were co-embedded with 20% artificial doublets to determine a threshold to distinguish doublets from singlets based on their nearest-neighbor network and a Doublet score was assigned based on each cell’s local neighborhood. For the multiome samples the RNA-doublet score was used as it proved slightly more stable. Additionally, the sex of the sample was determined based on the number of fraction of Y-chromosomal reads (>0.05% for male) as well TSS fraction. Nuclei that were not doublets, had more than 5,000 and less than 100.000 fragments, had more than 20% TSS fragments, had more than 1.000 RNA UMIs and at least 10% unspliced RNA UMIs were pooled to generate the main dataset (the final two filters only apply to multiome).

On average 27,599 high quality fragments per cell were identified with a Fragments in Peaks ratio of 54%. High quality nuclei were selected based on the number of fragments and fraction of fragments overlapping TSS as well as UMI count and splice ratio in multiome samples.

### Preclustering and consensus peak calling

The feature set used to generate the cell-by-peak matrices is dynamically derived from the data through peak calling. To do so the 20kb matrices are first joined and binarized after which the top 20% of autosomal bins are selected with an upper threshold of 60% coverage across the dataset and decomposed using Latent Semantic Indexing (LSI, more detailed description below). A KNN graph can then be constructed, and the data is clustered into broad clusters using Louvain clustering. Fragments from the nuclei belonging to each cluster are then aggregated and randomly split in two to generate two pseudo-bulk replicates per cluster. The pseudobulk aggregates are then down sampled to 25 million fragments and MACS2^55^ is used to call peaks using the following parameters: callpeak -f BEDPE -g hs –nomodel --shift 100 --ext 200 --qval 5e-2 -B –SPMR. Peaks were then extended to 400 bp using BEDtools and non-overlapping peaks between the pseudo-replicates were discarded. Next the identified peaks for all clusters were pooled and clustered using BEDtools cluster. For each cluster of peaks, the center point was extracted and extended to 400 bp to generate the consensus peak set. Peaks overlapping with the ENCODE blacklist were removed and the remainder was annotated using HOMER^56^ based on Gencode v32, after which the cell-by-peak matrix is generated.

### Latent Semantic Indexing

Decomposition was performed in two steps. First the matrix was depth normalized and infrequent features were upweighted by performing a Term-Frequency Inverse-Document-Frequency (TF-IDF) transformation. The resulting non-binary matrix was then used to compute the principal components using an incremental PCA. Initially 40 components are computed, but components that are not distributed significantly different from their predecessor are discarded along with a depth-correlated component if present. Next the components are batch corrected using Harmony to mediate chemistry and sample effects^12^.

### Clustering, embedding and aggregation

The cell-by-peak matrix is decomposed using an iterative LSI, meaning that the data is decomposed and clustered in two rounds. First the top 20.000 features by total coverage from the autosomal chromosomes are used to do a preclustering, after which 20.000 autosomal features are selected again based on the variance of their precluster level enrichment for a second LSI. Batch effects were again corrected for using Harmony. The second LSI is then used to generate nearest neighbor graphs and perform Louvain clustering. A tSNE is then generated using an adapted version of ‘the art of using tSNE’^57^ which better preserves global structure than native tSNE. Additionally, a UMAP is generated using UMAP-learn^58^ on default setting. For both methods Euclidean distances were used as a metric. Next all clusters were aggregated and a normalized Counts per Million (CPM) layer was added.

The enrichment of individual peaks was calculated as a pearson residual^59^. Briefly, fragments are modeled as a negative binomial distribution where the expected accessibility is the product of the total number of fragments per cluster and the fraction of fragments per peak. The residuals can then be calculated as the difference between the observed (Χ) and expected 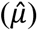 accessibility corrected by the negative binomial variation (dispersion parameter fixed at 100 for all analysis in this paper). For each cluster the top 2.000 peaks by pearson residual were marked as marker peaks. The 20.000 peaks with most variance between pearson residuals were used to calculate cluster similarities and to generate the cluster dendrogram.

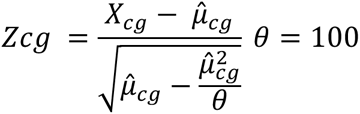

### Gene expression imputation & marker selection

31% of nuclei in the dataset were processed using the Single Cell Multiome ATAC + Gene expression kit. This allows for the imputation of gene expression measurements in the ATAC only samples. To predict gene expression in the scATAC-seq nuclei, first all multiome nuclei were all scaled to 5,000 UMIs and an ‘anchor’ net was first generated consisting of a directed graph of each scATAC-seq cell and their 10 nearest multiome neighbors. Next the weights were scaled to sum to 1 for each cell and the nearest neighbor matrix was multiplied with the gene expression profiles of the multiome nuclei to generate predicted gene expression profiles for each scATAC-seq cell.

Trinarization scores and gene enrichment were calculated as defined previously in^2^ with marker genes being selected based on their enrichment. The trinarization scores were then used for auto-annotation using a set of punch cards specific to early human development^4,5^.

### Subset analysis and pooling

The dataset was split based on cluster level marker expression into the following partitions: Fibroblast (*DCN*^+^ *COL1A1*^+^), Immune (*PTPRC*^+^), Vascular (*TAGLN*^+^ or *CLDN5*^+^ *FLT1*^+^), OPC (*PDGFRA*^+^ *OLIG1*^+^), RGL/GBL (*HES1*^+^ or *BCAN*^+^ *TNC*^+^) and Neuronal lineage (any of *INA*^+^ *NHLH1*^+^ *GAD2*^+^ *SLC17A6*^+^ *SLC6A5*^+^). The subsets were reanalyzed using the same pipeline described above. Clusters that contained less than 10 multiome nuclei or of which less than 1% of the total cluster size were multiome nuclei were excluded as well as clear clusters of doublets. The Neuronal lineage partition was split for a second round into the GABAergic lineage (*GAD2*^+^), Glutamatergic lineage (*SLC17A6*^+^, *SLC17A7*^+^ or *SLC17A8*^+^) and Peptidergic lineages. All partitions were then pooled again and new summary statistics and embeddings were generated.

### Motif enrichment

For every cluster the 2.000 selected marker peaks were used as input to HOMER findMotifsGenome^56^ using GC-matched genomic sequences as background. The Hocomoco v11 Full collection was chosen as the transcription factor binding motifs to be tested. The naming convention was manually altered to reflect genes names in the gene expression analysis. This allowed the filtering of false positives by exclusion of cell-motif combinations for which the corresponding transcription factor was unlikely to be expressed (trinarization score < .5). Additionally, all transcription factors were assigned to a family based on their archemotif^20^.

### Gene accessibility and cCREs

Gene accessibility scores were computed using an adapted version of the cicero workflow^14^ using the python SKGGM package. First, the distance parameter was estimated by optimizing the calculation of the regularized covariance matrices for 100 random 500 kb regions. Next the distance adjusted covariance for each accessible region with each TSS site was calculated in 500 kb bins with a 250 kb overlap. Most pairs are sampled twice and pairs with inconsistent covariances being discarded (∼5%). The co-accessibility cut-off was set empirically by testing the number of subnetworks over varying cut-off thresholds. Gene activity scores were then calculated by multiplying the peak-by-cell and region-to-TSS covariance matrices, normalizing against size factors derived from a linear regression model and pooling across the 25 nearest neighbors. Similarly to the region-TSS covariance matrix, cCREs were identified by calculating the region-Gene expression covariance.

### Identification of total accessible regions by sample

To identify general trends in opening/closing of chromatin, all fragments from individual cell classes and biological samples were pooled together and MACS2 was used to call peaks per class per sample. A one-sided fisher exact test with Benjamini-Hochberg correction was used to identify differential regions. A generalized linear model was used to estimate the influence of age on the number of accessible regions.

### VISTA enhancer overlap

CNS enhancers (from the VISTA database) were downloaded and lifted over to GRCh38 using UCSC liftOver, excluding any that could not be confidently lifted over, resulting in 620 enhancers, of which 596 overlapped with our peak set. The enhancers that were specific to the Forebrain, Midbrain and Hindbrain were isolated (total of 159, 75 and 78 respectively), similarly the corresponding peaks in the dataset were identified and the brain region with the highest accessibility was identified, after which the Jaccard similarity was calculated.

### Topic modelling

The full dataset was downsampled to a maximum of 10,000 nuclei per cluster to reduce computational burden and prevent overrepresentation. The number of topics was varied from 25 to 500 at intervals of 25, running for 50 iterations with an alpha of 50 divided by the number of topics and a beta of 0.1. The most stable model (175 topics) was selected based on topic-coherence and log-likelihood in the last iteration. The region-topic scores were normalized so that they summed to 1 for every cell and a tSNE-embedding was generated for the regions and binarized topic lists were generated by assigning each region to the topic that it scored the highest on. Next each topic was used as input for Homer2 with the Hocomoco transcription factors and the results were reduced to the highest scoring representative of each archemotif group. The binarized topics were also used as input for GREAT analysis^60^ to identify GO-terms describing each topic. For some selected terms the associated regions (within the topic) were used to calculate an enrichment score using pyCistopic’s signature_enrichment function.

### Enhancer Convolutional Neural Network

Nuclei from all clusters annotated as ‘Purkinje’, ‘Midbrain GABA’, ‘Cerebellum GNP’, ‘Hindbrain Glutamatergic’ or ‘Telencephalic Glutamatergic’ were grouped into 5 superclusters and enrichment between the clusters was recalculated and peaks were only included for learning if the log-fold change with the second highest accessibility was more than 1. Onehot encoded sequences (401 bp) were used as input to a Convolutional Neural Network trained as a classification model. The network consists of 4 convolutional layers of 256, 60, 60, 120 nodes and kernel sizes 7, 3, 5, 3 respectively, each layer was followed by batch normalization, RELU activation and maximum pooling. There were then two dense layers of 256 nodes with batch normalization, RELU activation and a dropout rate of 0.4. A softmax normalization was applied to the final output layer and cross entropy loss was used as the loss function with label smoothing set to 0.1. The model was trained using an Adam optimizer with a learning rate of 0.01. The model was trained for 26 epochs.

Contribution scores for each sequence were calculated using DeepLiftShap’s attribute function using the mean of the input sequence shuffled 100 times as background. The hypothetical score was calculated using for each possible nucleotide in the sequence by multiplying the contribution with the background corrected input^24^. TF-MoDisCo was then applied to all the sequences enriched in a cluster with a flanking size of 5 bp, a sliding window of 15 bp and a minimum cluster size of 30 seqlets.

### Pseudotime, GAMs and ChromVAR

For analysis of the Purkinje lineage all clusters labeled ‘Purkinje’ and the PTF1A^+^ cluster of Ventricular Zone Progenitors were isolated and a new TSNE was generated. PySlingshot was then used to calculate the pseudotime. pyGAM was used to fit gene and cCRE trends to the Purkinje Neuron lineage with gene expression being modeled using a PoissonGAM and cCRE accessibility using a LinearGAM. ChromVAR was applied using the JASPAR human PWM (human_pwms_v2) to compute motif variability.

### Supervised inference and stochastic simulation of Purkinje GRN

We used DELAY^25^ (https://github.com/calebclayreagor/DELAY) to infer the Purkinje GRN from gene-accessibility dynamics in pseudotime then performed stochastic simulations to verify the putative network’s gene-expression dynamics. First, we re-trained DELAY on a large scATAC-seq dataset of plasma B-cell differentiation^61^ with ChIP-seq ground-truth data^62^ to prepare the neural network to infer the Purkinje GRN from tens of thousands of single nuclei. Then, we fine-tuned DELAY on the Purkinje developmental trajectory using ground-truth targets of a cerebellar ataxia-related gene, ataxin-7^63^. For the final GRN inference, we used the expression-linked, log-normalized gene-linked peak counts from all TFs that were differentially expressed in at least 1% of Purkinje cells across pseudotime. We then used BoolODE^64^ to simulate the expression of each gene in the network given its top 8 most-likely regulators.

### GWAS enrichment

Accessible region locations were lifted over to GRCh37. Features were binarized on the cluster level with a Pearson Residuals threshold of 10. Cluster heritability was calculated using LD-score regression^41^. As a background we used the merger of our feature set with the features from development^10^ and adulthood^42^. Only SNPs from hapmap3 were included to reduce imputation errors. In total we tested 325 phenotypes from the UK Biobank^29^ and 11 psychiatric phenotypes ^30–40^. All used UK Biobank phenotypes had non-zero heritability estimates (Z-score > 4). Results UK Biobank phenotype enrichments were corrected for the number of cell types using FDR or Benjamini-Hochberg. For the psychiatric enrichments FDR and Bonferroni corrections were applied for the number of cell types and tests (α = 3.37e-5, 135 x 11 tests).

Two different MAGMA tests were conducted with default settings. First the cCRE linked to genes were annotated to genes in a custom MAGMA annotation file. A MAGMA gene analysis was used to assess which genes were affected in MDD. Next MAGMA gene analyses were conducted for ADHD, anorexia, ASD, MDD and schizophrenia, on a custom annotation filewhere individual accessible regions were treated like individual genes to identify specific deregulated elements. Accessible regions passing Benjamin-Hochberg correction were then used as input for Homer with the full vertebrate motif reference.

## Acknowledgments

We are grateful to the women who have decided to donate to science, making this study possible. We would also like to acknowledge the Developmental Tissue Bank (Department of Neurobiology, Care Sciences and Society, Karolinska Institutet) core facility, Human Developmental Biology Resource (United Kingdom) and Cambridge University for providing prenatal tissue samples. This research was supported by the NIHR Cambridge Biomedical Research Centre (BRC-1215-20014). The views expressed are those of the authors and not necessarily those of the NIHR or the Department of Health and Social Care. This work is funded by the following grants: Erling-Persson Foundation HDCA grant (SL), Knut and Alice Wallenberg Foundation grants 2015.0041, 2018.0172, 2018.0220 (SL), Swedish Foundation for Strategic Research SB16-0065 (SL) and the EU Horizon2020 BRAINTIME project 874606 (SL). We also acknowledge support from the National Genomics Infrastructure in Stockholm funded by Science for Life Laboratory, the Knut and Alice Wallenberg Foundation and the Swedish Research Council. We are grateful to Yang Eric Li and Bing Ren for hosting the online dataset. We thank Johannes Bastiaan Munting for providing supplementary code used while training the CNN. Finally, we thank Marek Bartosovic (Stockholm University), Mukund Kabbe (Karolinska Institutet) and Jens Hjerling-Leffler (Karolinska Institutet) for their relevant discussions.

## Rights to retention

For the purpose of open access, the author has applied a Creative Commons Attribution (CC BY) licence to any Author Accepted Manuscript version arising from this submission.

## Author contributions

C.M., S.L. designed the experiments in this study. C.M., L.H. performed experiments. C.M. analyzed and visualized the data. P.L. contributed to bioinformatics analysis.

C.R. analyzed Purkinje neuron transcription factor network. M.S., D.P. performed stratified GWAS. X.L., X.H., R.B., E.S. contributed to the sample collection. C.M. wrote the manuscript.

## Extended figures

**Extended figure 1:**
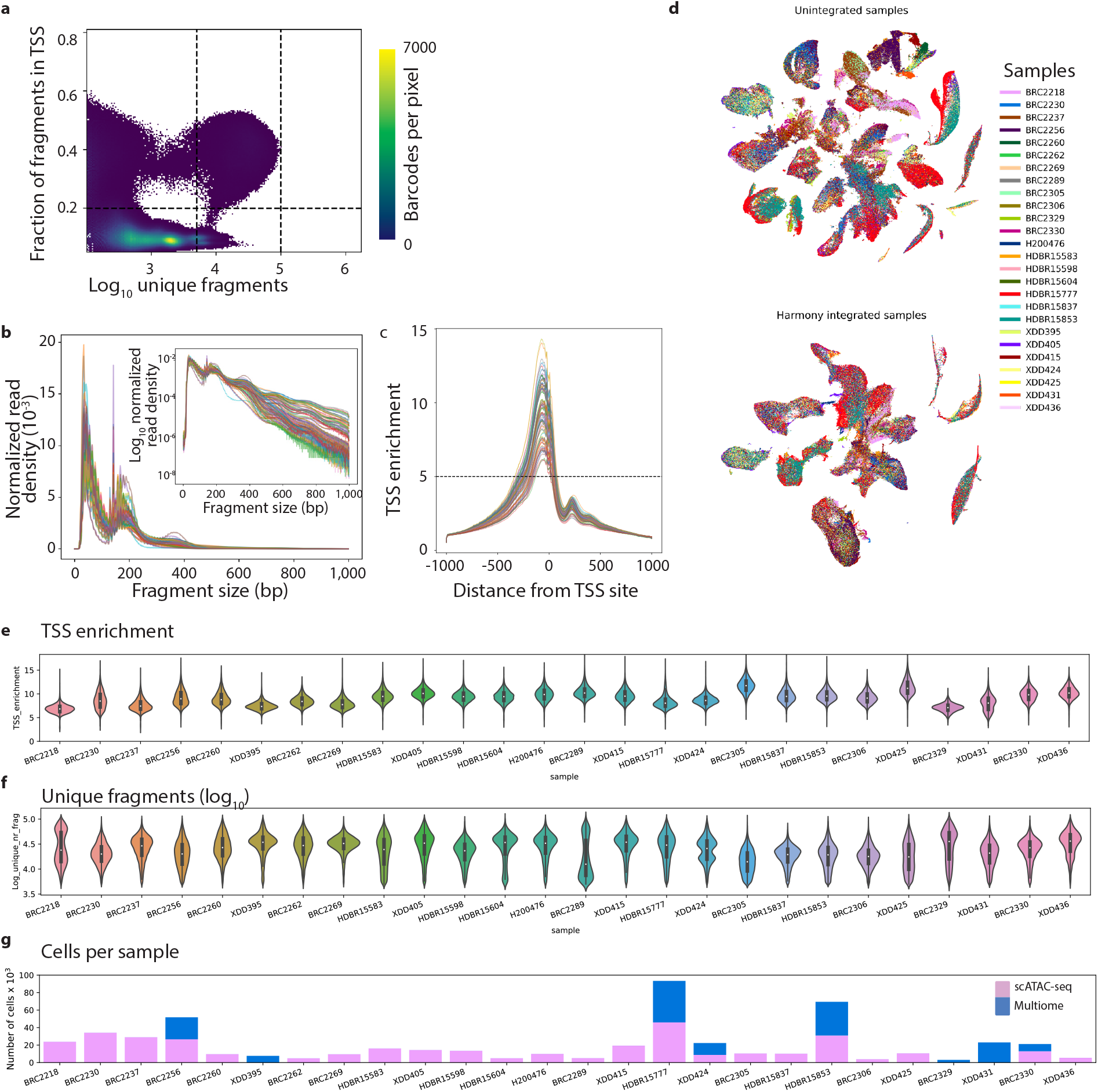
Quality control. A) Distribution of fragment count (log10) and fraction of fragments in TSS for collected barcodes. B) fragment size distribution per sample. Top plot shows log scaled density. C) t-SNE embedding generated from Latent Semantic Indexing without Harmony sample correction (top) and with sample correction (bottom). D) tSNEs of dataset without Harmony sample integration and with integration. Nuclei colored by sample ID. E) Distribution of TSS enrichment across nuclei per sample. 5 was used as a minimum sample level cut-off. F) Fragment count (log10) across nuclei per sample. G) Number of nuclei collected per sample, separated by method.

**Extended figure 2:**
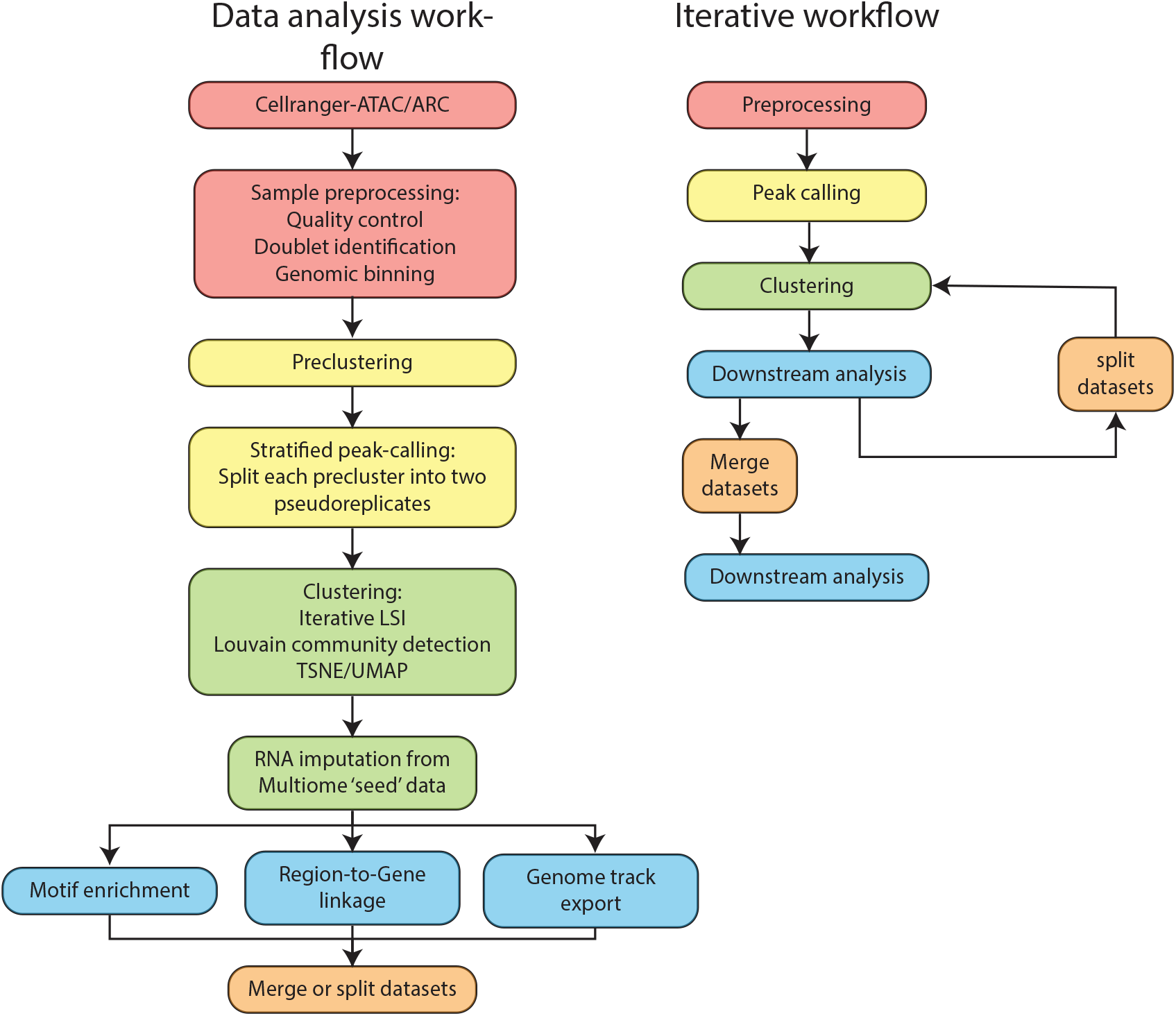
Analysis pipeline. Steps taken to analyze single cell data. Following quality control the dataset is clustered using genomic bins as features. Peak calling is then performed per cluster and a cell-by-peak matrix is generated and nuclei are clustered. The available multiome nuclei are then used to impute gene expression across the dataset. Downstream analysis is performed including motif enrichment analysis and region-to-gene linkage before splitting the dataset by cell class. Each subset is reclustered and reanalyzed separately before being pooled together again using the subset clusters and a final analysis round is conducted.

**Extended figure 3:**
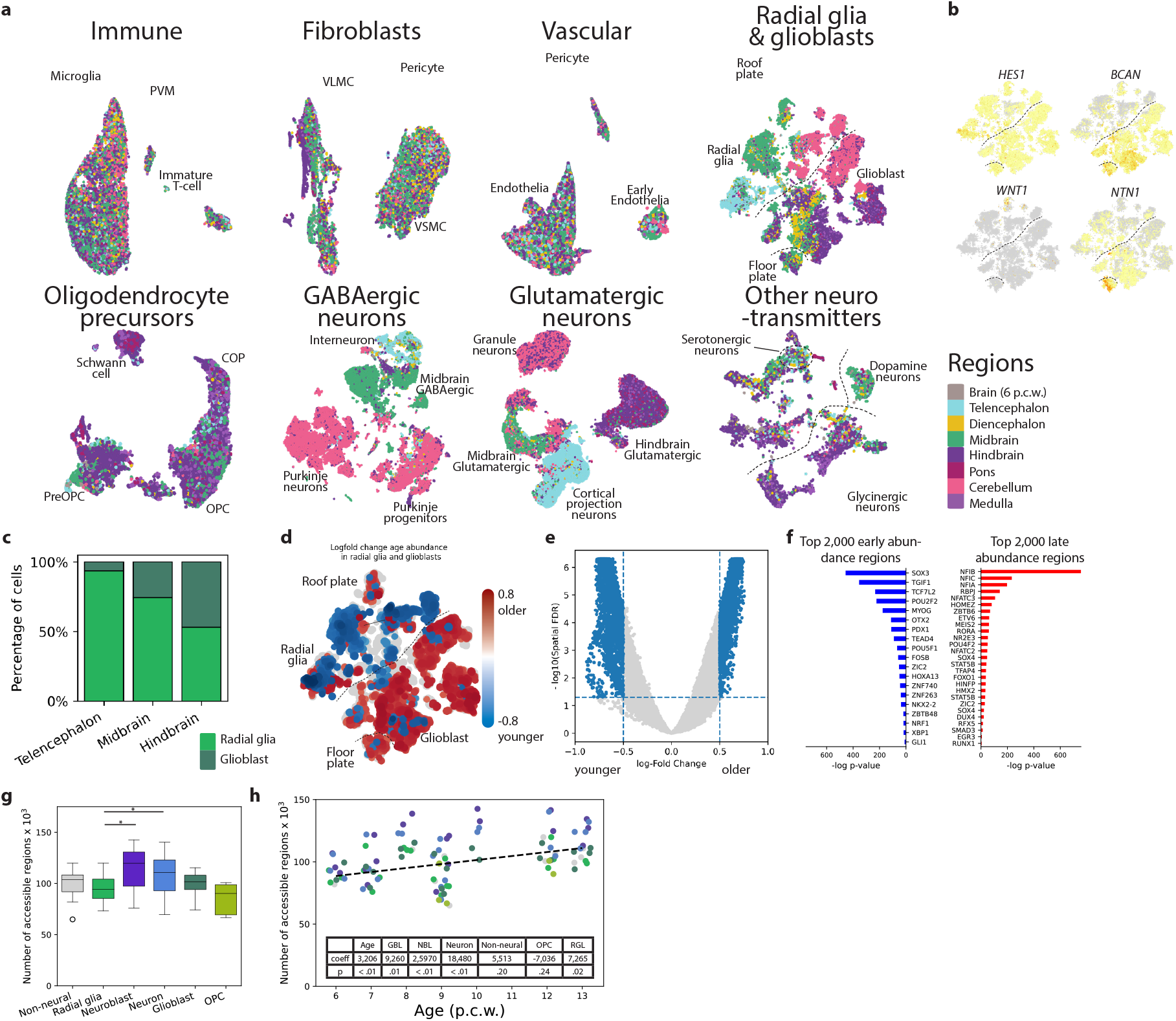
Regionalization of cell types. A) Annotation of region of origin for each cell class. While clear effects of regionalization can be seen in the neural lineage, the non-neuronal nuclei are more similar across brain regions. B) Expression of canonical markers used to annotate radial glia, glioblasts, the roof plate and the floor plate. C) Distribution of radial glia and glioblasts between brain regions. D) tSNE showing change in abundance of early vs. late nuclei in local neighborhoods among radial glia and glioblasts. E) Volcano plot of change in abundance of early vs. late nuclei in local neighborhoods among radial glia and glioblasts as identified using milopy. F) Enriched transcription factor motifs among the 2,000 most enriched accessible regions between early and late neighborhoods. G) Boxplot showing the number of accessible regions identified in different cell types. Neuroblasts and neurons have significantly more accessible regions than radial glia (two-sided independent t-test; neuroblast: t = 3.6; CI = 8,343-30,358; Cohen’s D = 1.13; 38 DF; p < 0.005; neurons: t = 2.5; CI: 2,485-22,706; Cohen’s D = 0.77; 43 DF; p < 0.05). H) Regression fitted to age to predict number of accessible regions using the cell types as covariates.

**Extended figure 4:**
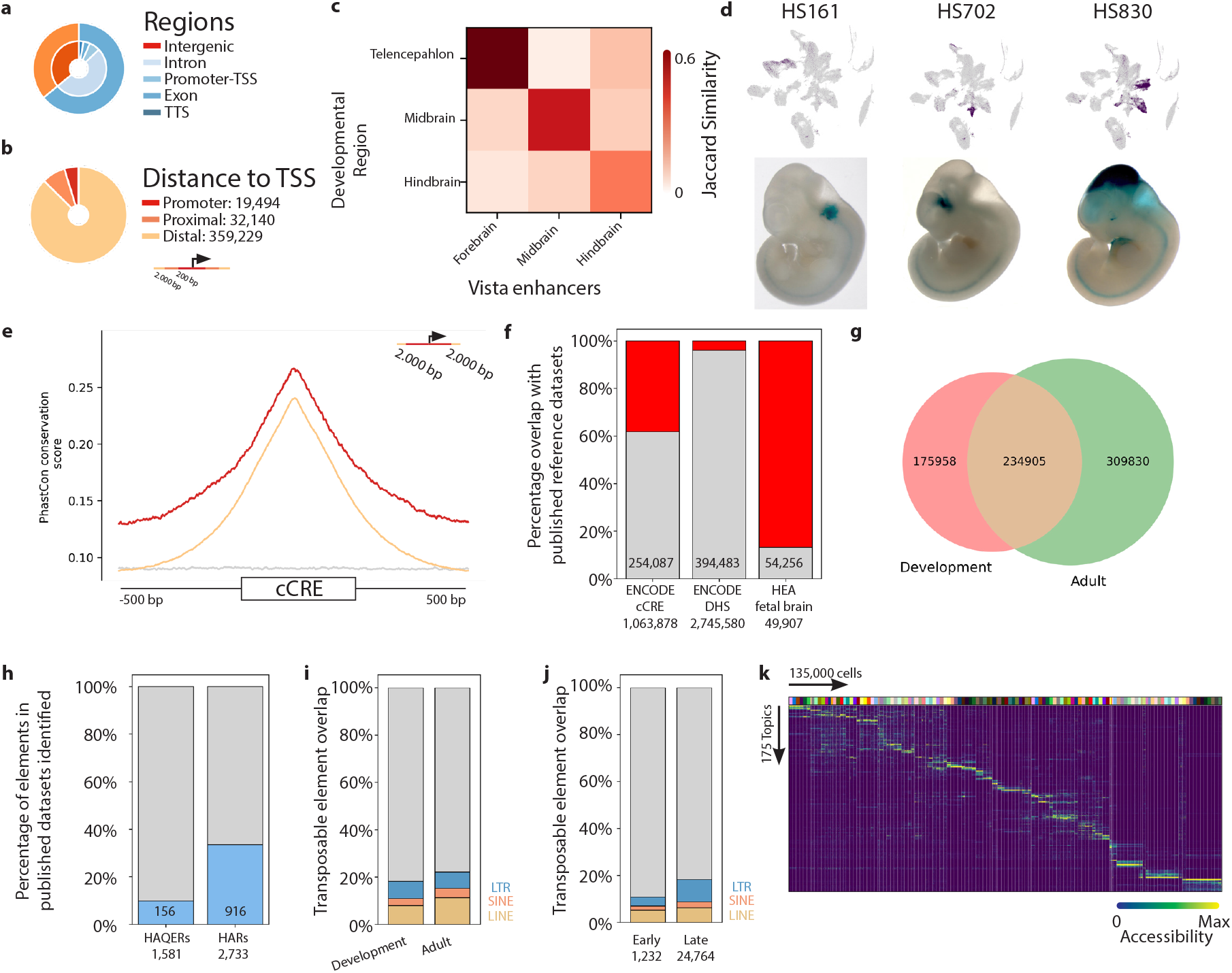
Annotation of accessible chromatin regions. A) Distribution of functional region annotations in relation to nearby genes. B) Distribution of accessible region distance to nearest TSS. C) Jaccard similarity between region-specific enhancers from the VISTA database and accessible regions identified in corresponding regions of the dataset. D) Spatially restricted accessibility of developmental enhancers overlapping with known enhancer sequences from the VISTA enhancer-database shown by LacZ staining. From left to right, active in Hindbrain neurons and glioblasts, immature interneurons in the Ganglionic Eminence and Midbrain radial glia and inhibitory neurons. E) Mean DNA conservation of proximal (<2,000 bp from TSS) and distal elements based on the PhastCon 100-way. F) Number of accessible regions that overlap with the ENCODE cCRE and DNAse hypersensitive site reference datasets. Additionally the number of elements that overlap with the human enhancer atlas fetal brain dataset. Red shows regions not in the reference dataset, gray are overlapping regions. G) Overlap between the identified accessible regions in this study (development) and a comparable study in the adult human brain (Li *et al.,* 2022 under revision). H) Overlap with two sets of evolutionarily accelerated regions, with overlap in blue and regions from the comparison list not in our dataset in gray. Human accelerated regions (HARs) are regions with increased rates of nucleotide substitution that are conserved in other species, while human ancestor quickly evolved regions (HAQERs) are regions that diverged rapidly between humans and chimpanzees that were not previously constrained. I) Overlap with annotated transposable elements. J) Comparison of transposable elements in early vs. late nuclei across the dataset. G) Heatmap of region topics across the 135,00 nuclei included in the topic modeling.

**Extended figure 5:**
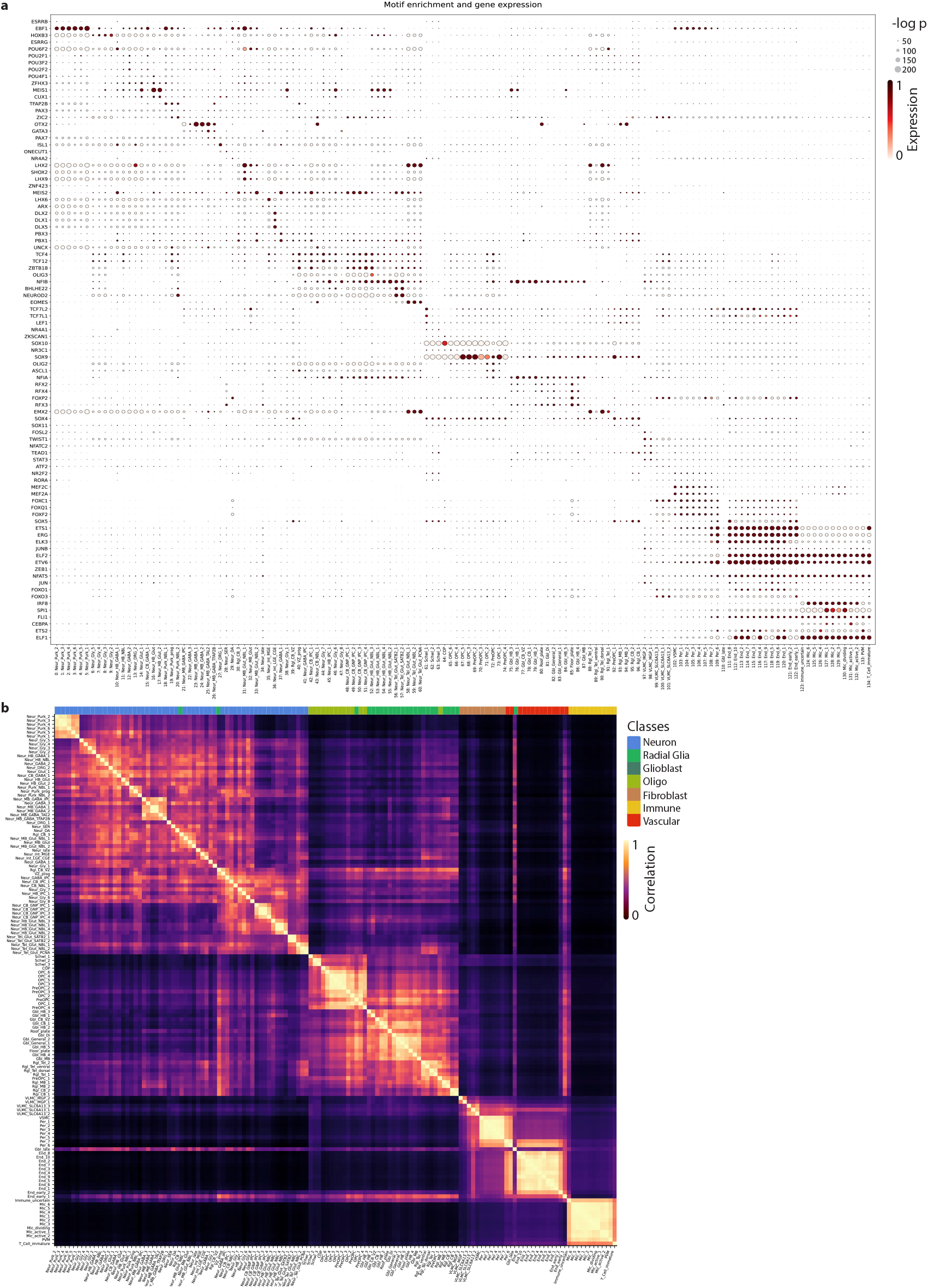
Cluster annotation. A) Extended motif enrichment plot of all clusters. Dot size represents -log p-value of the motif enrichment. The color represents expression level. B) Correlation between cell types based on marker peak accessibility. The colorbar at the top represents the assigned cell class.

**Extended figure 6:**
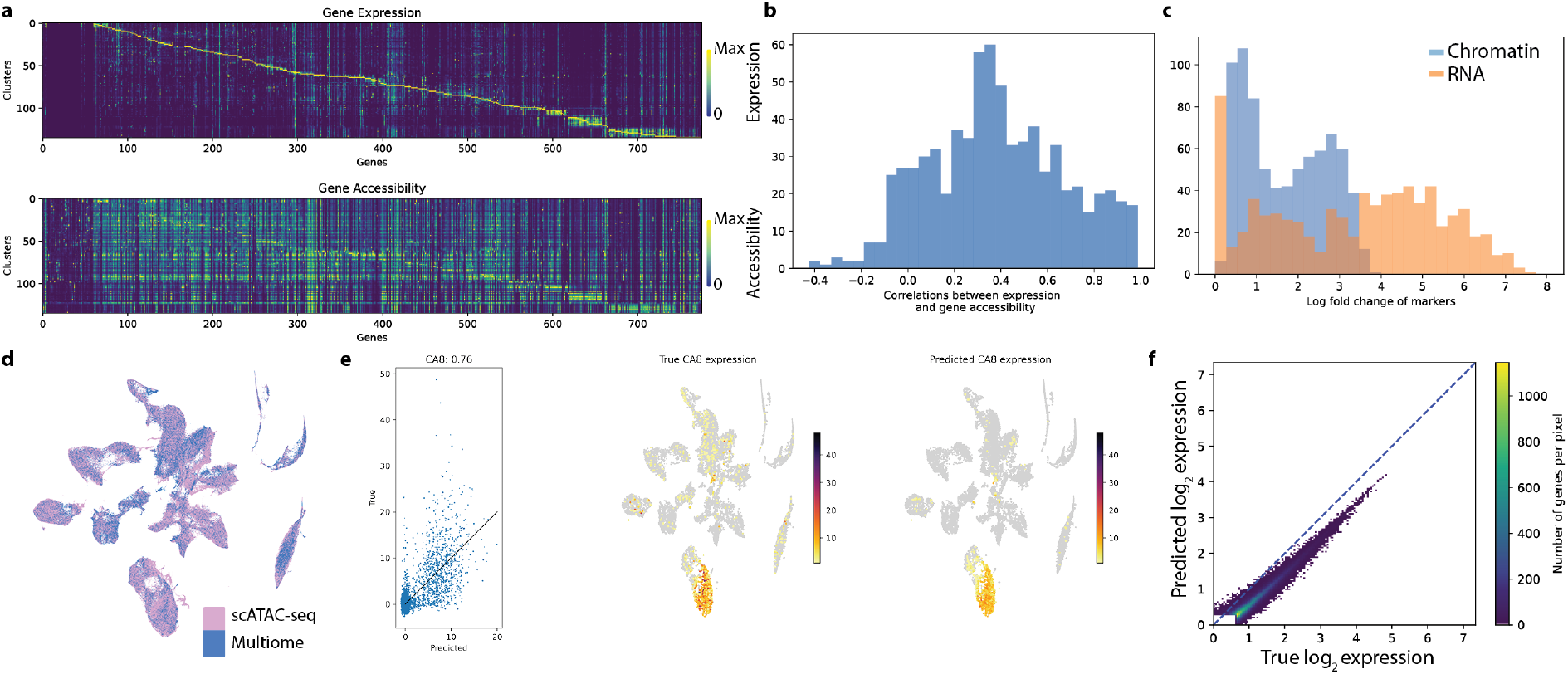
Gene expression imputation. A) Comparison between cell type specificity of marker genes in gene expression and gene accessibility space. Top 5 enriched genes were selected for each cluster in both modalities and the union was used for plotting. B) Correlation between expression and accessibility of each selected marker gene. C) Maximum log fold change of marker versus median expression. The genes with RNA enrichment of zero are not expressed. D) Distribution of scATAC-seq and multiomic nuclei across the tSNE embedding. E) Leave-one-out validation of imputed expression. Predicted versus true expression of *CA8*. F) Top 100,000 gene-cluster expression pairs true vs imputed expression.

**Extended figure 7:**
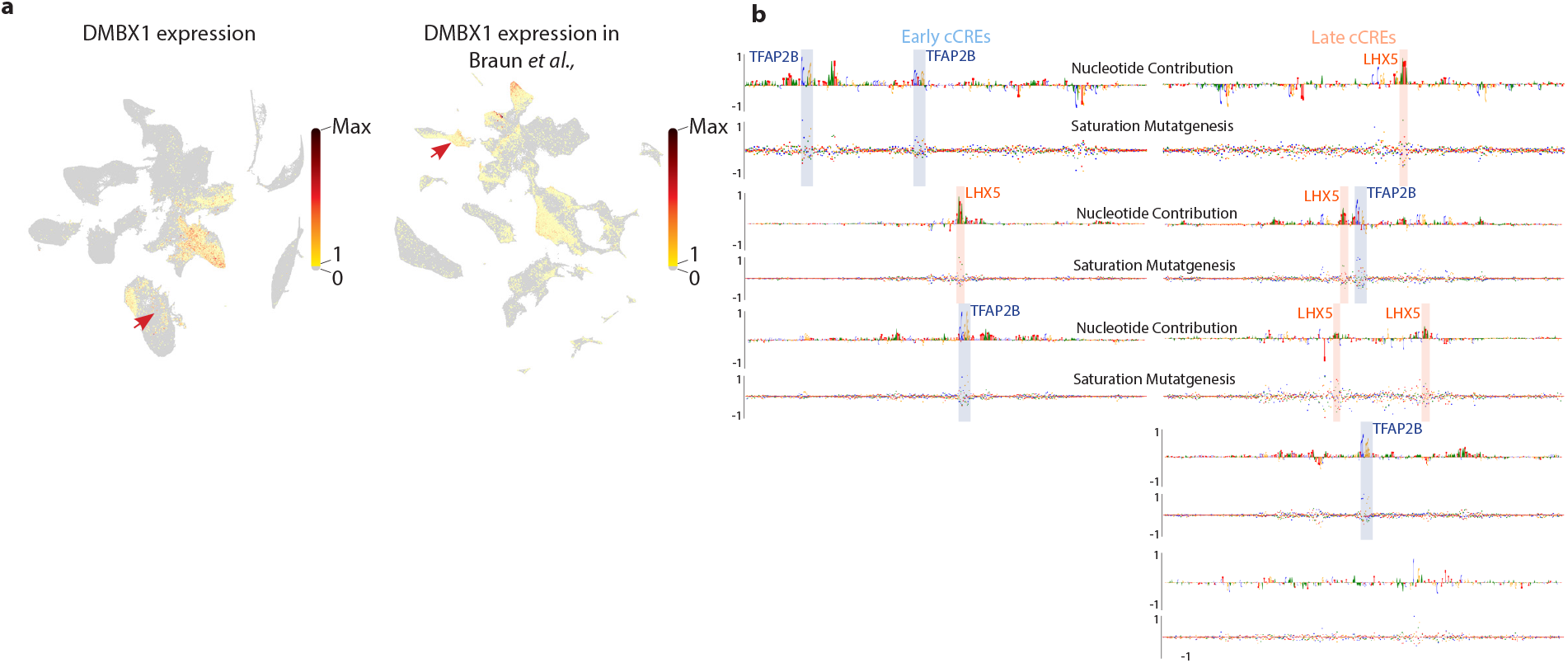
Additional figures related to TF CNN model. A) Expression of *DMBX1* in this dataset and Braun *et al.,*^5^ B) Additional cCREs upstream of *ESRRB*.

**Extended figure 8:**
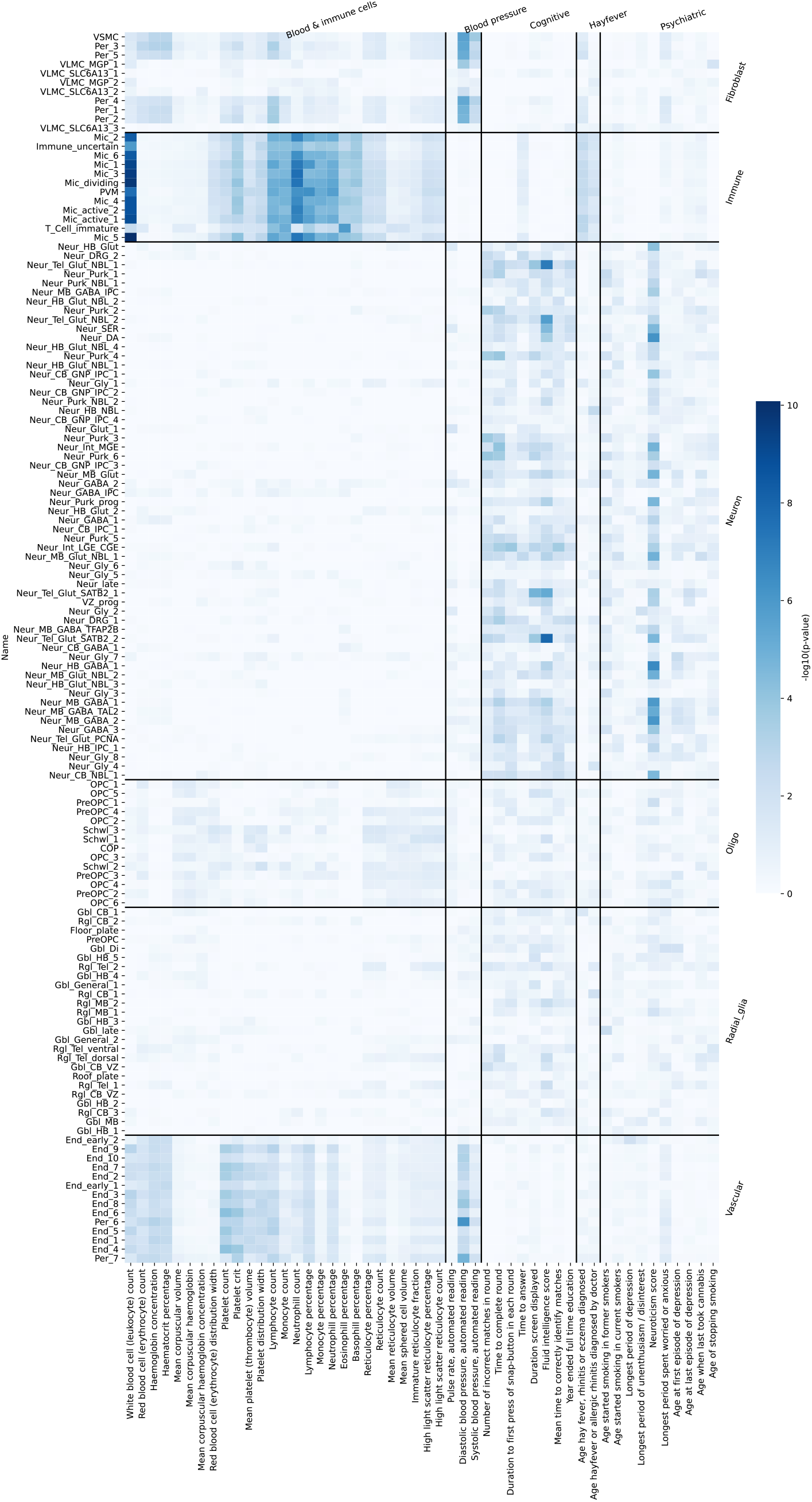
Enrichment of selected UK Biobank traits in neurodevelopmental cell types. Each trait is assigned to a group of related traits (Immune, blood pressure, cognitive, hayfever and psychiatric). Clusters are ordered by major cell class. Most traits show the expected enrichment pattern.

**Extended figure 9:**
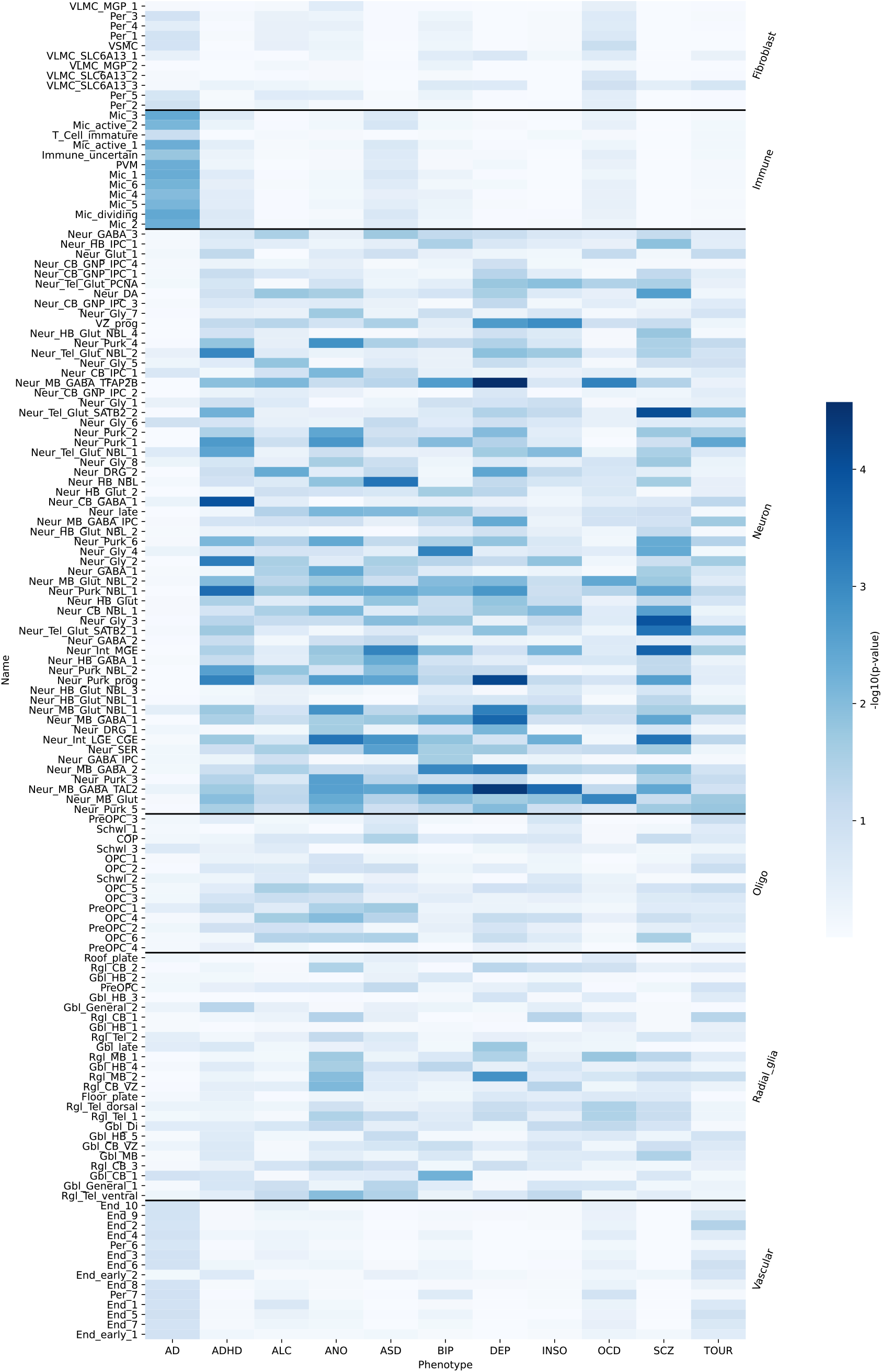
Enrichment of selected psychiatric phenotypes in neurodevelopmental cell types. Clusters are ordered by major cell class. While not reaching significance after multiple test correction, an increased association between Alzheimer’s disease and immune cells can be observed in opposition to the other traits which primarily are associated with neuronal cells.

